# Pulsed Field Ablation Enables Stromal Manipulation and Tumor Reduction in Pancreatic Ductal Adenocarcinoma

**DOI:** 10.64898/2026.10.06.756887

**Authors:** MacKenzie V. Demmel, Lincoln N. Strickland, Kyle McAndrews, Khadija Turabi, Alyssa Waller, Shwetapadma Dash, Urvinder Kaur Sardarni, Nirav C. Thosani, Raju Viswanathan, Jennifer M. Bailey-Lundberg

## Abstract

Pancreatic ductal adenocarcinoma (PDAC) is characterized by a dense desmoplastic microenvironment that promotes therapeutic resistance and disease progression. Here, we investigated the impact of pulsed field ablation (PFA), a non-thermal electroporation-based therapy, on tumor and stromal remodeling in a syngeneic murine PDAC model. PFA induced rapid apoptotic and necrotic tumor cell death, transient neutrophil infiltration, and durable suppression of tumor growth. Single-cell RNA sequencing following repeated PFA treatments revealed enrichment of a classical secretory tumor cell state associated with complement and stress-response pathways and reduced epithelial-to-mesenchymal transition (EMT) programs. Although overall immune composition was largely unchanged, PFA markedly remodeled the stromal compartment, decreasing αSMA-positive activation and shifting cancer-associated fibroblasts (CAFs) from tumor-promoting myofibroblastic CAFs toward *Dpp4*+ complement-secreting CAFs. This transition was accompanied by increased IL-1 signaling potential, reduced *Tgfb1* expression, and diminished tumor cell TGF-β response signatures. Serial PFA treatments further suppressed EMT-associated programs, identifying stromal reprogramming as a potential mechanism through which PFA enhances tumor control.

**Statement of Significance:** PFA suppresses PDAC growth while reprogramming the tumor stroma. PFA shifts CAFs from myofibroblastic to *Dpp4+* complement-secreting states, reduces TGF-β and EMT signaling, and promotes durable tumor control through stromal remodeling.

## INTRODUCTION

Pancreatic ductal adenocarcinoma (PDAC) remains one of the deadliest malignancies despite recent therapeutic advances. A defining feature of PDAC is its extensive desmoplastic stroma, which comprises the majority of tumor mass and contributes to therapeutic resistance, immune exclusion, and disease progression^1-4^. Within this tumor microenvironment, cancer-associated fibroblasts (CAFs) regulate extracellular matrix deposition, immune cell recruitment, and tumor cell behavior through complex paracrine signaling networks^5,6^. Recent studies have revealed substantial CAF heterogeneity in PDAC, with distinct fibroblast populations exerting divergent effects on tumor progression and therapeutic response^3,7-12^. Understanding how emerging treatment modalities influence these stromal populations may therefore provide new opportunities to improve patient outcomes.

Local tumor ablation has emerged as an attractive strategy for PDAC treatment because it can provide direct tumor control while minimizing systemic toxicity^13-19^. Conventional techniques, including radiofrequency ablation, microwave ablation, and cryoablation, rely on hypothermic or thermal injury to induce cell death^13,15,20-22^. While effective at destroying tumor tissue, thermal approaches may damage surrounding structures and can promote tissue remodeling responses that reinforce stromal barriers^17^. Consequently, there is increasing interest in non-thermal ablative technologies capable of achieving tumor destruction while preserving the surrounding microenvironment.

Pulsed field ablation (PFA) is a non-thermal ablation modality that uses high-intensity electrical pulses to induce irreversible electroporation and cell death. Since its mechanism is independent of thermal injury, PFA can selectively ablate tumor tissue while largely preserving vasculature and nearby critical structures^23^. Although PFA is being increasingly investigated in solid tumors^24^, the biological consequences of PFA extend beyond direct tumor cell killing and remain incompletely understood, including how PFA treatment influences stromal composition and CAF phenotypes. Given the central role of CAFs in regulating immune responses, extracellular matrix organization, and tumor cell plasticity, understanding these effects may reveal previously unrecognized mechanisms of therapeutic benefit^2,3,9-11,25-27^. Here, we use histologic, immunologic, and single-cell transcriptomic analyses to characterize the acute and long-term effects of PFA on the PDAC tumor microenvironment. We demonstrate that PFA treatment induces extensive tumor destruction while simultaneously remodeling stromal cell states, promoting a shift from tumor-promoting myofibroblastic CAFs toward tumor-restraining *Dpp4*+ complement-secreting CAFs (csCAFs). This transition is associated with suppression of TGFβ-responsive and epithelial-to-mesenchymal transition-associated tumor programs, suggesting that PFA may alter not only tumor viability but also the biological state of the tumor microenvironment.

## METHODS

### Preclinical KPC Subcutaneous Tumor Model

All mouse model procedures followed UNMC’s IACUC protocol and adhere to ARRIVE guidelines. 500,000 KPC cells were prepared in a PBS:Matrigel mix (1:1) and injected subcutaneously in both flanks of 8-week-old male C57BL/6 mice; however, for this study, only the effects of PFA treatment on the treated tumors were analyzed. KPC cells were derived in the Tuveson Lab from *Kras*^*LSL-G12D/+*^*;Trp53*^*LSL-R172H/+*^*;Pdx1-Cre* mice, which develop PDAC. Sex as a biological variable: since the KPC cell line was derived from male mice, these experiments used male mice as they were found to produce the most consistent growth rates. Tumors were measured three times per week with a vernier caliper. Tumor volume was calculated as (length × width × width)/2 in mm^3^. Growth rate was calculated by taking the linear regression of all tumor volume measurements before treatment and all tumor measurements after treatment. The day of treatment was the final tumor volume for the before-treatment growth rate and the first tumor volume for the after-treatment growth rate. Statistical analysis was performed using a two-way ANOVA or a mixed-effects analysis in Prism GraphPad software.

### Pulsed Field Ablation

Prior to treatment with the ablation or sham, mice were anesthetized with isoflurane via a nose cone. Skin around the ablation site was cleaned with 3 alternating rounds of 70% ethanol and iodine and a 2.5 mg/kg dose of bupivacaine (NDC 0409-1162-18, Hospira Inc.) and 5 mg/kg dose of meloxicam (NDC 13965-559-20, Vet One) was administered through subcutaneous injection prior to ablation. A custom PFA bipolar needle probe and generator system manufactured by ALPFA Medical (Palo Alto, California) was used to deliver ablation. The ALPFA PFA probe was inserted for a total of 3 ablations per session. Mice were observed for signs of pain post-ablation. The sham control mice were transferred to the procedure room, placed under anesthesia, received bupivacaine and meloxicam, the probe was inserted, but no ablation was performed. **Figure 1 Protocol:** ablation was performed 18 days post tumor implantation. An incision was made in the center of the tumor, and both prongs of the ALPFA PFA probe were inserted. The tumor on the right flank of the mouse was ablated. **Figure 2 Protocol:** ablation was performed 18 days post tumor implantation. An incision was made in the center of the tumor, and both prongs of the ALPFA PFA probe were inserted. The larger tumor was ablated. **Figure 3A Protocol:** ablation was performed 9, 16, and 23 days post tumor implantation. Prongs of the ALPFA PFA probe were placed on either side of the tumor. The tumor on the right flank of the mouse was ablated. **Figure 3E Protocol:** ablation was performed 15, 18, and 21 days post tumor implantation. One prong of the ALPFA PFA probe was inserted in the tumor. The tumor on the right flank of the mouse was ablated.

**Figure 1.**
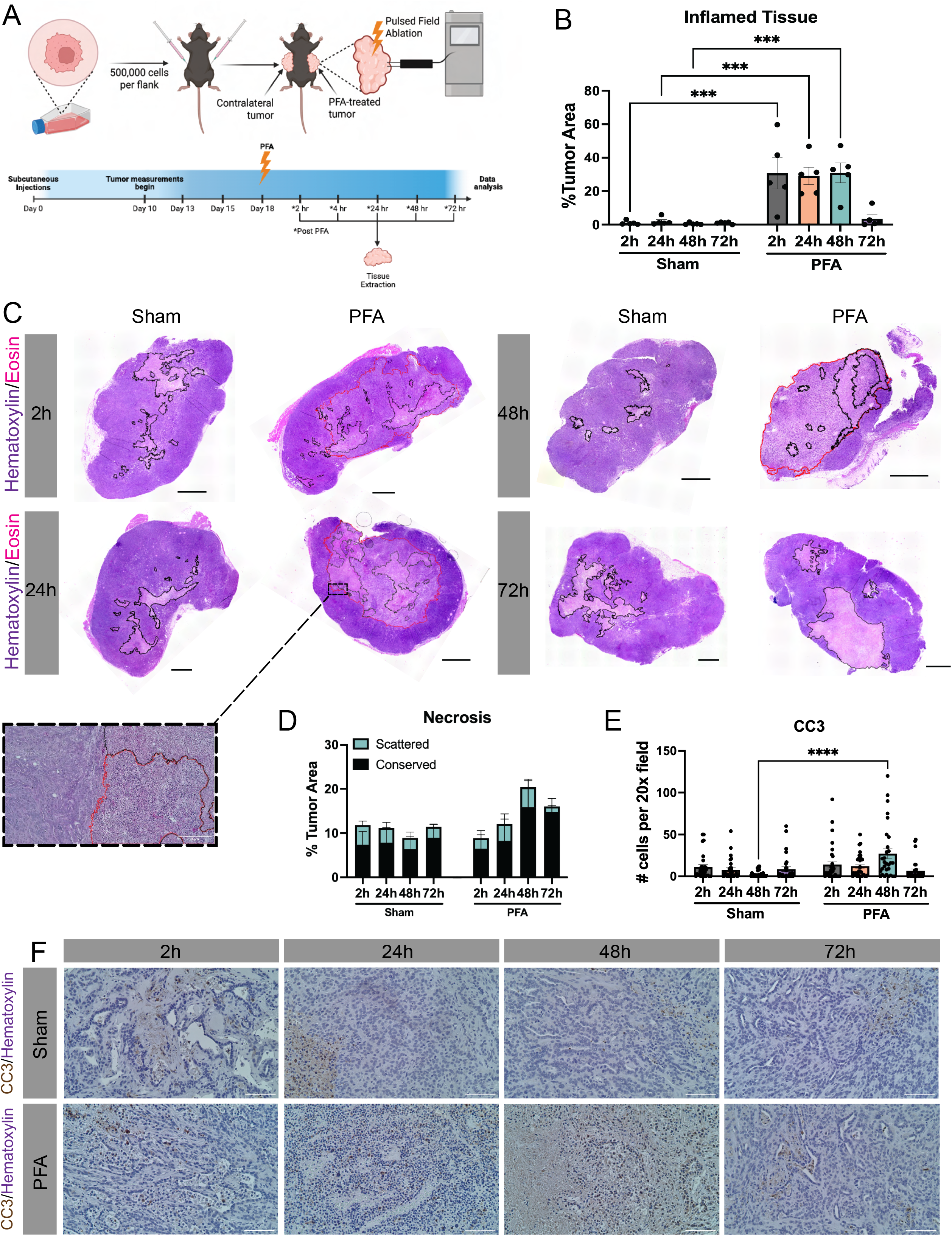
Bipolar PFA produces acute tumor destruction. **A.** Experimental design of PFA-treated mice. Protocol was duplicated in Sham-treated mice with probe insertion but no ablation (n=5). **B**. Percentage of total tumor area that displayed an inflamed phenotype. Inflamed tissue area was significantly higher in PFA tumors up to 48 hours post ablation. **C**. Full-slide H&E staining at 2, 24, 48, and 72 hour time-points post ablation or sham. Inflamed tissue/ablated region circled in red; necrosis circled in black. **D**. Percentage of total tumor area that had either conserved necrosis (area > 2mm^3^) or scattered necrosis. **E**. PFA had significantly more CC3+ cells than sham at 48 hours post ablation (n=3, 10 images per tissue). **F**. Representative CC3 IHC images at each time point. Data presented as mean ± SEM. \**p<0.05.**p<0.01, ***p<0.001, ****p<0.0001*.

**Figure 2.**
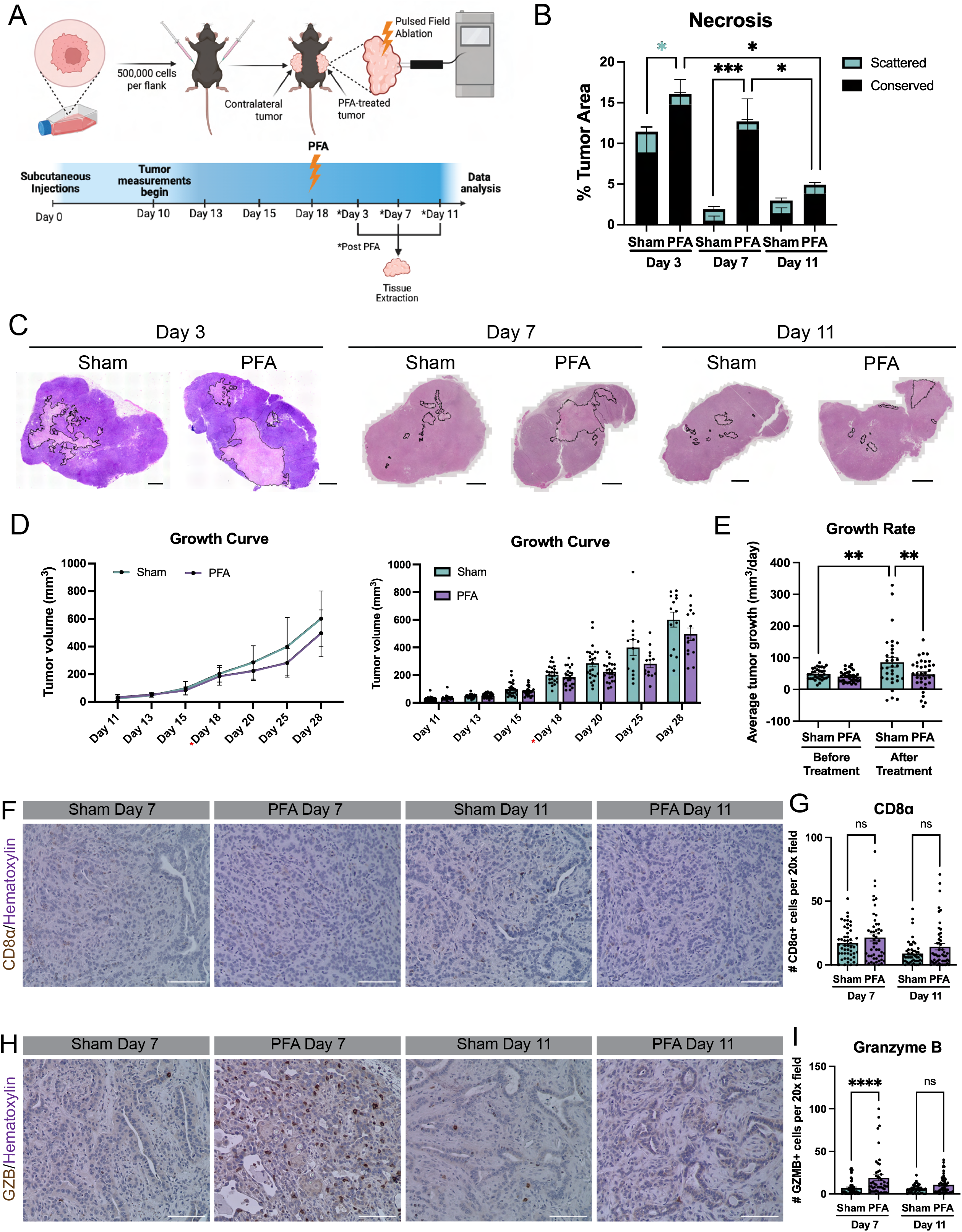
PFA reduces growth rate of PDAC tumors *in vivo* and alters the tumor microenvironment. **A.** Experimental design of PFA-treated mice. Protocol was duplicated in Sham-treated mice with probe insertion but no ablation (Day 3: n=5, Day 7: n=11, Day 11: n=14). **B**. PFA induces sustained necrosis up to 7 days post ablation (Day 3: n=5, Days 7 and 11: n=10). **C**. Full-slide H&E staining at 3, 7, and 11 days post ablation or sham. Necrosis circled in black. **D**. Growth curves for PFA-treated and Sham-treated tumors. Day of ablation marked by asterisk. **E**. PFA significantly reduces growth rate compared to Sham treatment. **F**. Representative IHC images of CD8⍰at each time point. **G**. There is no significant difference between CD8⍰levels at either time point (n=5, 10 images per tissue). **H**. Representative IHC images of GZB at each time point. **G**. There is significantly more GZB in the PFA tumors at 7 days post ablation when compared to sham (n=5, 10 images per tissue). Data presented as mean ± SEM. \**p<0.05. **p<0.01*,*\*\*\*p<0.001, ****p<0.0001*.

**Figure 3.**
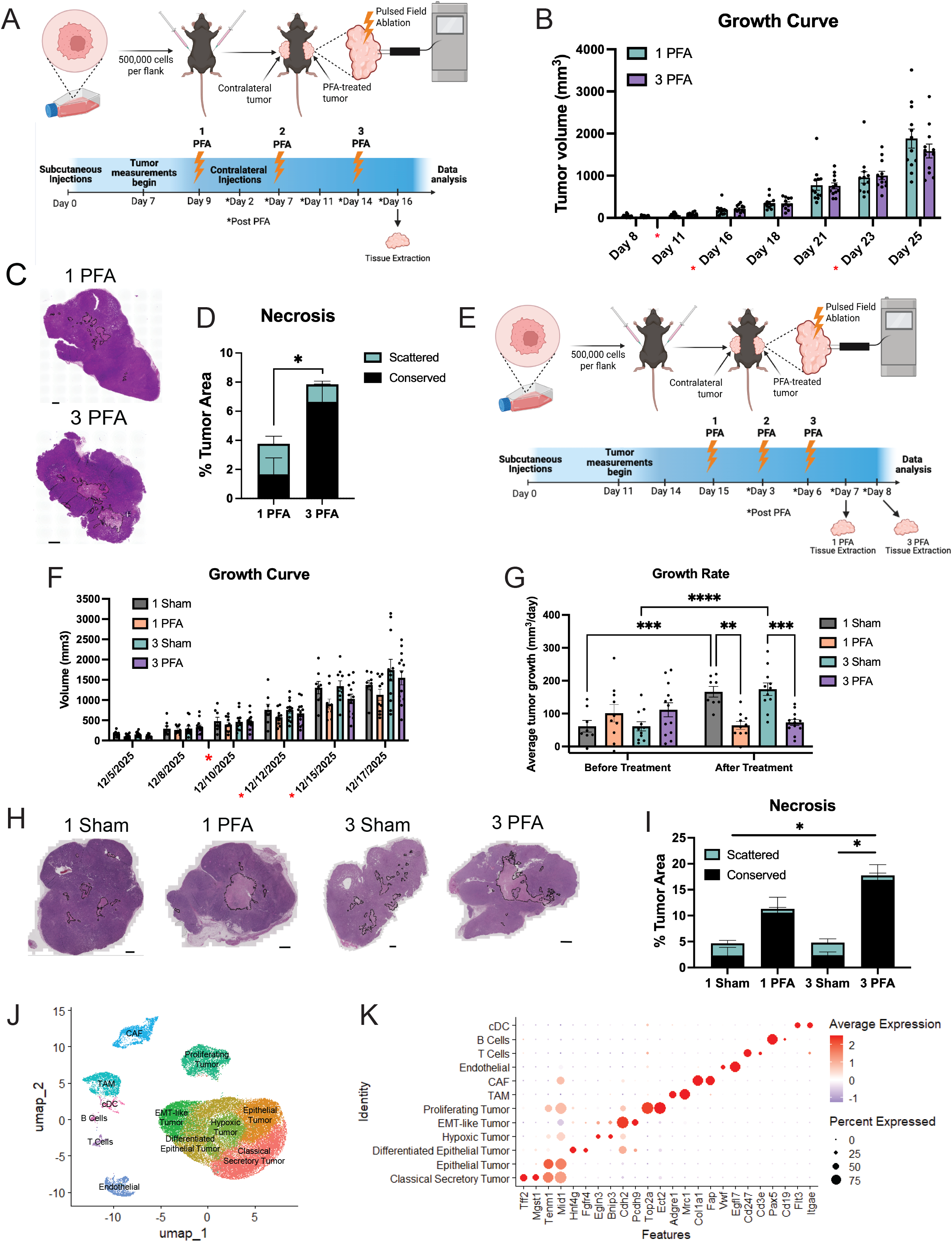
Multiple PFA sessions increases necrosis and reduces growth rate. **A.** Experimental design of PFA-treated mice. Protocol was duplicated in Sham-treated mice with probe insertion but no ablation (1 PFA: n=12, 3 PFA: n=11). **B**. Growth curves for PFA-treated and Sham-treated tumors. Days of ablation marked by asterisk. **C**. Full-slide H&E staining for each treatment group. Necrosis circled in black. Scale bars 1 mm. **D**. Necrosis is significantly increased in 3 PFA tumors. **E**. Experimental design of PFA-treated mice. Protocol was duplicated in Sham-treated mice with probe insertion but no ablation (1 Sham: n=8, 1 PFA: n=10, 3 Sham: n=11, 3 PFA: n=13). **F**. Growth curves for PFA-treated and Sham-treated tumors. Days of ablation marked by asterisk. **G**. 1 PFA session and 3 PFA sessions both significantly decrease tumor growth rate when compared to 1 Sham and 3 Sham. **H**. Full-slide H&E staining for each treatment group. Necrosis circled in black. Scale bars 1 mm. **I**. 3 PFA sessions significantly increases necrosis compared to 1 sham and 3 sham. **J**. Blinded UMAP embedding of 9 tumors (n=3 for 3 Sham, 1 PFA, and 3 PFA) with labeled clusters. **K**. Top genes used to identify the cell type corresponding to each cluster. Data presented as mean ± SEM. \**p<0.05. **p<0.01*,*\*\*\*p<0.001*,*\*\*\*\*p<0.0001*.

### Hematoxylin and Eosin Staining and Quantification

Hematoxylin and eosin (H&E) staining was performed as previously described^17^. Statistical analysis was performed using a two-way ANOVA in Prism GraphPad software.

### Immunohistochemistry and Quantification

Immunohistochemistry (IHC) staining and quantification was performed as previously described^18,28^. Statistical analysis was performed using a two-way ANOVA in Prism GraphPad software. Antibodies and their corresponding dilutions are in **Supplemental Table 1**.

### Single cell RNA-seq and Analysis

Tumors were cut in half along the length and half of the tumor was flash frozen. The flash-frozen half for three samples per group were sent to NovoGene and processed. The RNA library was aligned to the GRCm39_2024-A reference genome. Raw sequencing data were processed using Cell Ranger (v.7.0.0), generating gene expression feature-barcode matrices. The unfiltered feature-barcode matrices were used as input for CellBender Remove Background (v0.2.0)^29^ to identify cell-associated barcodes and remove ambient RNA contamination. CellBender was run on the following parameters: 16,000 expected cells, 30,000 total droplets, false positive rate of 0.01, and 150 epochs. Data was then processed using Seurat package (v5.4.0)^30^. To ensure data quality, cells were excluded if they fell outside of 200 to 8,000 features, 200 to 50,000 RNA counts, or had lower than 20% mitochondrial reads. Raw counts were normalized with the function *NormalizeData*. Doublet detection was performed using scDblFinder (v1.24.10)^31^. Principal component analysis (PCA) was then performed on the 3,000 highly variable genes. Samples were integrated using harmony (v2.0.5)^32^. The *FindNeighbors* function was used with the first 30 PCs to build the nearest neighbor graph and the *FindClusters* function was used with a 0.3 resolution to identify cell clusters. The UMAP method was employed for dimensionality reduction and 2D visualization of cell clusters. Differential expression analysis was performed using the *FindMarkers* function with Wilcoxon ranked-sum test. One cluster was found to be dominated primarily by ribosomal genes and was determined to be a low-quality artifact. Distribution of the cluster across samples and a list of the top differentially expressed genes are provided in **Supplemental Fig. 1**. The cluster was subsetted out, and the remaining cells were re-clustered into their final state. A table reporting the number of cells retained after each QC step is provided in **Supplemental Table 2**, with a total retention of 79% and 132,791 total cells. Cell types were annotated by examining the expression of canonical cell markers in cell clusters. The difference in gene expression between two conditions of a cell type was tested using Wilcoxon ranked-sum test and adjusted using the Benjamini-Hochberg procedure. A table of the packages used in the analysis and formation of graphs is provided in **Supplemental Table 3**^**33-38**^.

### Data Availability Statement

The sequencing data generated in this study have been deposited in the Gene Expression Omnibus (GEO) and will be made publicly available upon publication.

## RESULTS

### Bipolar PFA produces acute tumor destruction

To characterize an immediate tumor response to PFA treatment, we performed one PFA or Sham session after allowing a subcutaneous tumor to grow for 18 days, then we extracted the tumor at 2 hours, 4 hours, 24 hours, 48 hours, and 72 hours post ablation **(Fig. 1A)**. From hematoxylin and eosin staining (H&E), we detected a phenotypically distinct region that we quantified as inflamed tissue. This region showed similar patterns with reported tumor responses to irreversible electroporation (IRE)^39^, with small, dense nuclei, areas of cellular drop out, and low cytoplasmic boundaries. The region resolved into either necrosis or viable tissue by 72 hours post ablation **(Fig. 1B-C)**. Since this region was specific to PFA-treated tumors, we called it the “ablated region.”

We also quantified necrosis and noticed it developed in a single region in PFA tumors rather than the scattered necrosis we observed in Sham-treated tumors. We distinguished these two types of necrosis as “conserved” if the area was greater than 2 mm^2^ or “scattered” if it was less. Conserved necrosis trended upwards at progressive time points in PFA when compared to Sham **(Fig. 1C-D)**. At 48 hours post ablation, there was significantly more cleaved caspase 3 (CC3) as quantified from IHC **(Fig. 1E-F)**. This corresponds with a significant infiltration of MPO+ neutrophils in the PFA-treated tumors at 48 hours post ablation, which were localized to the ablated region. These neutrophils were no longer present by 72 hours post ablation **(Supplemental Fig. 2A-C, 2F)**. There was no observable increase in CD8⍰ or Granzyme B (GZB) in PFA at these time points, although both stainings also demonstrated localization within the ablated region **(Supplemental Fig. 2D-F)**.

### PFA reduces growth rate of PDAC tumors in vivo and alters the tumor microenvironment

To determine the long-term effects of PFA treatment in PDAC, we performed a single ablation session and extracted tumors at 7 days and 11 days post ablation. The results were compared to 3 days post ablation from the previous experiment to examine the acute and long-term effects **(Fig. 2A)**. While there was high necrosis in the Sham-treated tumors on Day 3, this necrosis was resolved by Day 7. However, PFA-treated tumors maintained a significantly high percentage of necrosis at the Day 7 time point. This necrosis was significantly reduced by 11 days post ablation **(Fig. 2B-C)**. The tumor volume remained consistently, but not significantly, lower in PFA-treated mice compared to Sham-treated mice even at 11 days post ablation **(Fig. 2D)**. Also, PFA-treated tumors grew significantly slower than Sham-treated tumors after treatment **(Fig. 2E)**. When comparing CD8⍰T cell infiltration, there was no significant difference between treatment as found by IHC **(Fig. 2F-G)**. However, there was significantly higher staining for GZB at Day 7 in the PFA-treated tumors, indicating that the T cells present could be more cytotoxic at that time point **(Fig. 2H-I)**.

### Multiple PFA sessions increase necrosis and reduce growth rate

Since necrosis was largely resolved by 11 days post ablation in PFA-treated tumors, we wanted to determine whether additional ablations spaced 7 days apart could re-introduce sustained cell death **(Fig. 3A)**. We found that tumors treated with a single ablation session (1 PFA) and three ablation sessions (3 PFA) maintained similar tumor volumes throughout **(Fig. 3B)**. However, the 3 PFA tumors had significantly more necrosis than the 1 PFA tumors **(Fig. 3C-D)**.

We then wanted to investigate whether we could induce sustained phenotypic changes to the TME consistent with what we observed at 48 hours post ablation. So, we performed an experiment with ablations spaced three days apart and extracted the tumors which received 3 PFA or Sham sessions 48 hours after the final session. Tumors which received 1 PFA or Sham session were extracted 7 days after the single ablation **(Fig. 3E)**. From this, we found that raw tumor volume was reduced in both 1 PFA and 3 PFA when compared to 1 Sham and 3 Sham respectively **(Fig. 3F)**. Similarly, 1 PFA and 3 PFA were found to significantly reduce tumor growth rate **(Fig. 3G)**. H&E staining showed an increase in necrosis in 1 PFA and 3 PFA, with 3 PFA significantly higher than 1 Sham and 3 Sham **(Fig. 3H-I)**. Three samples each from the groups 3 Sham, 1 PFA, and 3 PFA were then processed for single cell RNA sequencing (scSEQ). From this, 12 distinct cell types were identified, including the following tumor, stromal, and immune cell clusters: B cells, CAFs, conventional Dendritic cells (cDCs), classical secretory tumor, differentiated epithelial tumor, EMT-like tumor, endothelial, epithelial tumor, hypoxic tumor, proliferating tumor, T cells, and tumor associated macrophage (TAM) clusters **(Fig. 3J)**. These cell types were classified by their top differentially expressed genes **(Fig. 3K)**.

### PFA enriches a classical secretory phenotype and alters stromal phenotype

Six tumor clusters were identified from the scSEQ data. Each cluster had a distinct genetic signature that corresponded to a classical secretory, epithelial, differentiated epithelial, epithelial-to-mesenchymal-transition (EMT)-like, hypoxic, and proliferating tumor subtype **(Fig. 4A)**. The tumor subtype proportions were largely consistent across groups **(Fig. 4B)**. However, the classical secretory subtype was elevated in 1 PFA tumors, and even more so in 3 PFA tumors **(Fig. 4C)**. We investigated this cluster further and found that it was elevated in the gene ontology pathways for regulation of opsonization and apoptosis signaling in response to endoplasmic reticulum (ER) stress **(Fig. 4D)**. It also scored significantly higher for a classical PDAC score^40^ and gene ontology plasma membrane repair score **(Supplemental Figure 3A)**. Gene set enrichment analysis (GSEA) identified the hallmark oxidative phosphorylation pathway and coagulation pathway enriched in this cluster, though the leading-edge genes associated with the coagulation pathway were primarily complement-associated **(Fig. 4E-4G)**. This cluster was thus determined to be stressed, but without EMT-associated aggression, and with the complement and opsonization pathway expression.

**Figure 4.**
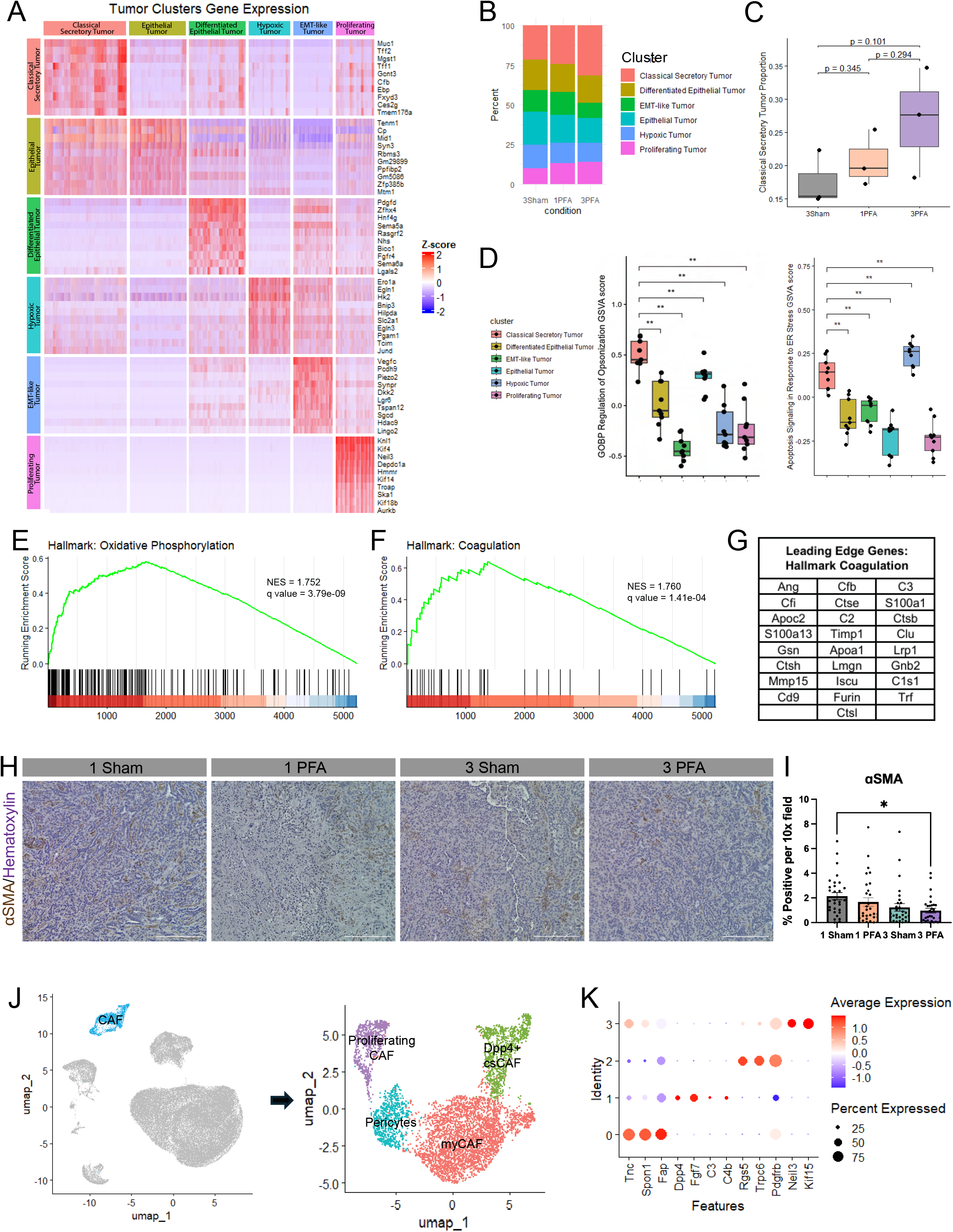
PFA enriches a classical secretory phenotype and alters stromal phenotype. **A.** Heatmap of the six tumor clusters with the top 10 differentially expressed genes in each cluster. **B**. Tumor cluster proportion across each condition.**C**. Classical secretory tumor proportion is slightly elevated in 1 PFA and more elevated in 3 PFA tumors. **D**. The classical secretory tumor cluster is enriched in GOBP regulation of opsonization and apoptosis signaling in response to ER stress pathways. **E**. The classical secretory tumor cluster is enriched in the Hallmark oxidative phosphorylation pathway. **F**. The classical secretory tumor cluster is enriched in the Hallmark oxidative phosphorylation pathway. **G**. Leading edge genes for the coagulation pathway in the classical secretory tumor cluster. **H**. Representative IHC images for αSMA. Scale bar 25μm. **I**. αSMA is significantly reduced in 3 PFA tumors compared to 1 Sham (n=3, 10 images per tissue). **J**. CAF sub-clustering reveals four distinct subtypes. **K**. Top differentially expressed markers for each CAF subtype. Data presented as mean ± SEM. *\*p<0.05. **p<0.01*.

We did not identify a difference in the proportion or subtypes of the immune populations present **(Supplemental Fig. 4A-O)**. There was also no change in NIMPR14 or MPO IHC staining between groups **(Supplemental Fig. 5C-5F)**. However, the stroma composition proved to be highly differential. While there was no significant difference in trichrome-identified collagen deposition **(Supplemental Fig. 5A-B)**, we found that alpha smooth muscle actin (αSMA) was significantly reduced in 3 PFA tumors **(Fig. 5H-I)**. αSMA did not show significant reduction at earlier time points following one PFA session compared to Sham treatment **(Supplemental Fig. 5G-H)**. To more deeply investigate these results, we analyzed the CAF population from scSEQ. After sub-clustering, four distinct populations were revealed **(Fig. 4J)**. These clusters were identified by their top differentially expressed markers and were found to correspond to a myCAF, *Dpp4*+ csCAF, pericytes, and proliferating CAF population **(Fig. 4K)**.

**Figure 5.**
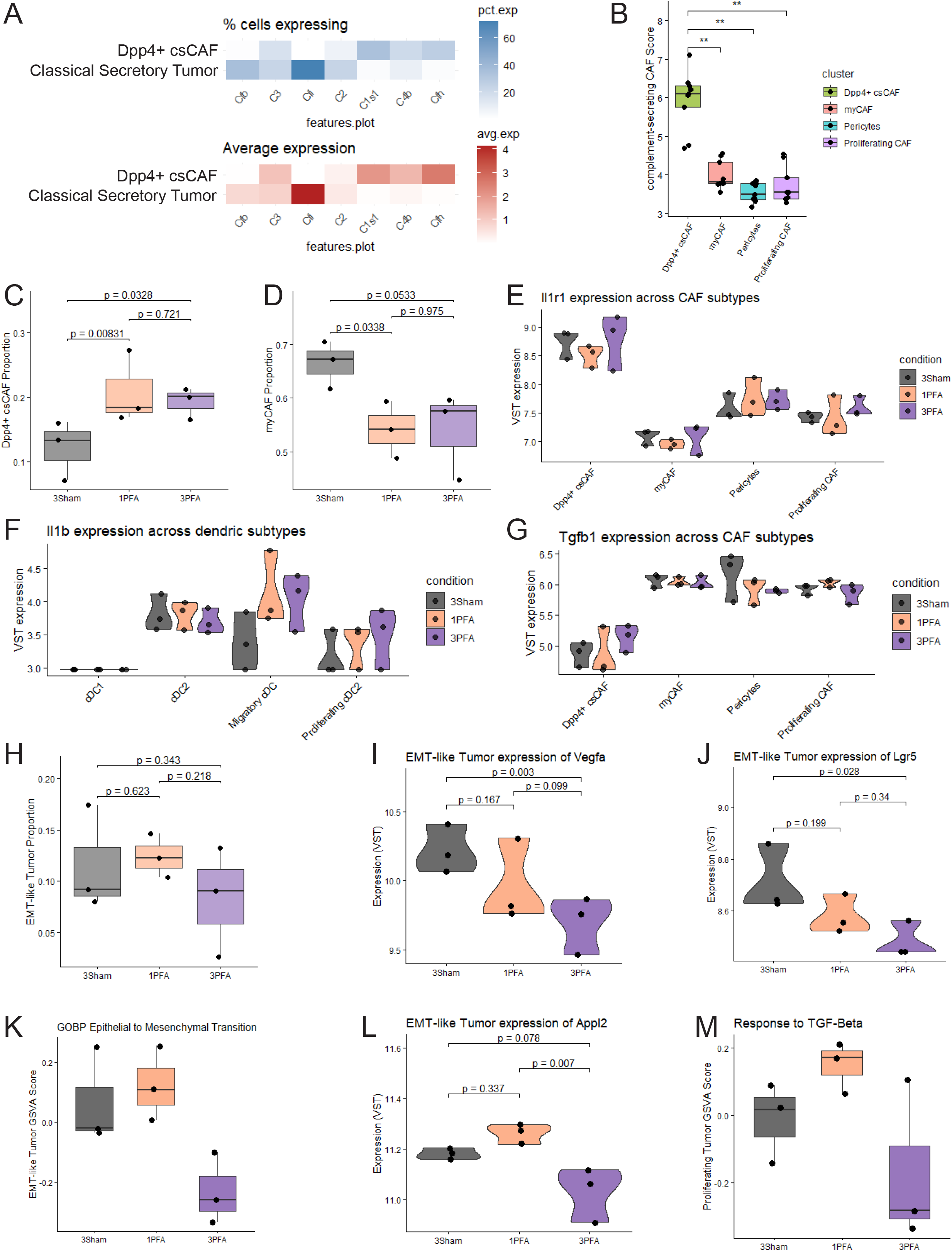
PFA reduces tumor aggression. **A.** Heatmap comparing percent and average expression of key complement-associated genes between the classical secretory tumor cluster and Dpp4+ csCAF cluster. **B**. Dpp4+ csCAFs score significantly higher for mouse csCAF associated genes than other CAF clusters. **C**. Dpp4+ csCAFs are significantly enriched in both 1 PFA and 3 PFA. **D**. myCAF proportion is significantly lower in both 1 PFA and 3 PFA. **E**. Variance-stabilized-transformed expression of IL-1 receptor *Il1r1* across CAF subtypes. **F**. Variance-stabilized-transformed expression of IL-1 producer *Il1b* across dendritic cell subtypes. **G**. Variance-stabilized transformation expression of *Tgfb1* across CAF subtypes. **H**. Proportion of EMT-like tumor cells is consistent between groups. **I**. Variance-stabilized-transformed expression of EMT-associated gene *Vegfa* is significantly lower in 3 PFA. **J**. Variance-stabilized-transformed expression of EMT-associated gene *Lgr5* is significantly lower in 3 PFA. **K**. 3 PFA has a reduced score for the epithelial to mesenchymal transition gene ontology pathway. **L**. Variance-stabilized-transformed expression of TGF-β-response-associated gene *Appl2* is significantly lower in 3 PFA. **M**. 3 PFA tumors have a reduced score for the response to TGF-β gene ontology pathway. Bar plots presented as mean ± SEM. *\*p<0.05. **p<0.01*.

### PFA reduces tumor aggression

As both a CAF and tumor subpopulation were found to express the complement pathway, we looked at the expression of complement genes between the clusters and found that expression was distributed between clusters **(Fig. 5A)**. Indeed, the *Dpp4*+ csCAFs scored significantly higher than the other CAF subtypes for genes associated with csCAFs in PDAC^41^ **(Fig. 5B)**. This population was significantly enriched in both 1 PFA and 3 PFA tumors, while the myCAF proportion was significantly lower **(Fig. 5C-D)**. Investigation of the potential mechanism causing this shift revealed that Dpp4+ csCAFs produced comparatively high amounts of the IL-1 receptor *Il1r1* **(Fig. 5E)**. This corresponded with *Il1b* expression in the dendritic cell (DC) population, specifically sub-clustered migratory DCs in 1 PFA and 3 PFA tumors **(Fig. 5F)**. Sub-clustering of the cDC cluster revealed 4 distinct dendritic cell clusters: conventional DC1, conventional DC2, proliferating cDC2, and migratory cDC clusters **(Supplemental Fig. 3B-C)**. While the DC populations were found to have different expression profiles across conditions, the proportion of subtypes was relatively consistent, excluding a slight shift from conventional DC1 to conventional DC2 in 1 PFA **(Supplemental Fig. 3D-G)**. Finally, we observed that the *Dpp4+* csCAFs expressed lower levels of *Tgfb1* **(Fig. 5G)** and higher levels of *Tgfbr3* **(Supplemental Fig. 3H)** than all other CAF subtypes.

To determine whether this shift in the stroma was altering the tumor phenotype, we looked deeper at the EMT-like tumor population. While the proportion was similar between groups **(Fig. 5H)**, we found a slight reduction in expression of EMT-associated genes *Vegfa* and *Lgr5* in 1 PFA with a significant reduction in 3 PFA tumors **(Fig. 5I-J)**. Consequently, when scored against the gene ontology epithelial-to-mesenchymal transition pathway, 3 PFA tumors showed a large reduction **(Fig. 5K)**. Similarly, we found that 3 PFA tumors had significantly lower expression of *Appl2*, a gene associated with TGF-β response **(Fig. 5L)** and scored lower in gene ontology’s response to TGF-β pathway **(Fig. 5M)**.

### PFA initiates sequence resulting in tumor death and lower tumor aggression

From these data, we have observed that PFA induces cell death, as observed by increased CC3 and necrosis. **(Fig. 1C-F, Fig. 2B-C, Fig. 3C-D, Fig 3H-I)**. Similarly, we observed enrichment in a tumor cluster associated with stress and opsonization **(Fig 4C-D)**. This corresponded with an influx of neutrophils at early time points **(Supplemental Fig. 2A-C)**. We also observed an increased expression of *Il1b* in the migratory dendritic cells in 1 PFA and 3 PFA tumors, which corresponded with a shift in CAF proportion towards *Dpp4+* csCAF: a CAF subset which has elevated levels of *Il1r1* **(Fig. 5C-F)**. Since the EMT-like tumor cluster scored lower in both response to TGF-β and epithelial to mesenchymal transition in 3 PFA tumors **(Fig. 5K, 5M)** we propose a mechanism wherein PFA-induced cell death leads to recruitment of neutrophils, clearance of opsonized cells, and a shift in the expression of the dendritic cells. This expression could therefore cause the shift from myCAF to *Dpp4*+ csCAF, reducing the TGF- β levels in the TME and thus reducing EMT-like aggression **(Fig. 6)**.

**Figure 6.**
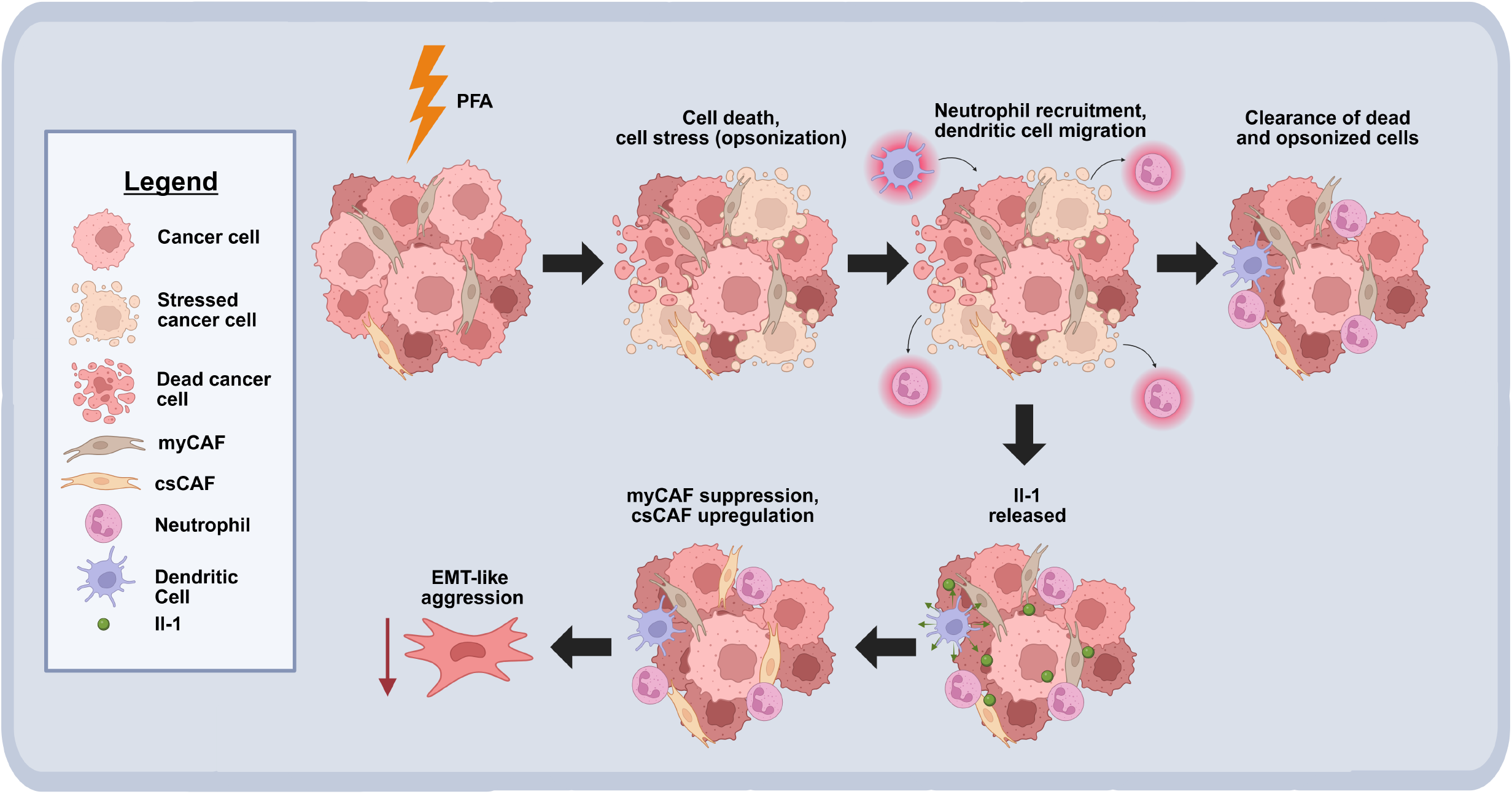
PFA initiates sequence resulting in tumor death and lower tumor aggression. Graphical abstract depicting proposed mechanism. PFA induces cell death, resulting in enrichment of a tumor cluster associated with stress and opsonization, corresponding with recruitment of neutrophils, clearance of opsonized cells, and a shift in the expression of the dendritic cells. Additionally, there is a shift in the CAF populations from myCAFs toward a *Dpp4+* csCAF with elevated levels of *Il1r1*, therefore reducing EMT-like aggression.

## DISCUSSION

Analysis of the early response to PFA treatment revealed rapid and spatially restricted tumor injury. Necrosis was largely confined to the central region of the tumor corresponding to the site of ablation, while increased CC3 staining was observed 48 hours after treatment, indicating ongoing cell death beyond the immediate ablation event. These histologic changes were accompanied by increased infiltration of MPO+ neutrophils, consistent with a role for innate immune cells in the clearance of necrotic debris and damaged tumor cells. In contrast, CD8α and GZB staining were not increased during the early post-ablation period. Instead, GZB levels were significantly reduced at 2 hours following treatment, suggesting acute loss of cytotoxic immune populations within the ablated region in addition to direct tumor cell killing.

By 24 hours, both CD8α and GZB staining had returned to levels comparable to sham-treated tumors, with positive cells localized primarily within the ablated area. Together, these findings reveal a rapid sequence of localized tissue injury, immune cell depletion, and early immune repopulation following PFA treatments.

To determine whether these responses were sustained after a single ablation session, tumors were evaluated 7 and 11 days after treatment. Although tumor growth remained significantly suppressed, the extent of necrosis declined between days 7 and 11, indicating progressive resolution of ablation-induced injury. Similarly, GZB staining was elevated at day 7 but diminished by day 11. These data suggest that the direct cytotoxic effects and associated immune responses triggered by a single PFA treatment are transient despite persistent suppression of tumor growth. Importantly, these findings define a temporal window during which the tumor microenvironment is actively undergoing remodeling following ablation.

We next investigated whether repeated PFA treatments could further alter the tumor microenvironment. Two ablations administered 7 days apart resulted in greater tumor necrosis than a single treatment, whereas three ablations delivered at 3-day intervals produced comparable effects on tumor volume and necrosis. Despite these similar bulk tumor responses, single-cell RNA sequencing demonstrated substantial differences in tumor and stromal cell states following three serial PFA treatments. Tumors exposed to three ablations were enriched for a stressed tumor-cell population and exhibited significantly reduced α-SMA staining, consistent with a decrease in myCAFs. In parallel, the proportion of csCAFs was increased. These observations suggest that three PFA treatments have effects that extend beyond direct tumor ablation and can induce substantial remodeling of stromal cell populations.

The mechanisms underlying these stromal changes remain to be fully defined. However, csCAFs expressed high levels of the IL-1 receptor, *Il1r1*, and IL-1 signaling has previously been implicated in regulating CAF state transitions in PDAC^42^. Consistent with this possibility, dendritic cells exhibited increased *IL-1* expression following both single and repeated PFA treatment sessions. While these data do not establish a causal relationship, they support a model in which PFA-induced inflammatory signaling contributes to the observed shift in CAF composition.

The biological consequences of this stromal remodeling may be significant. Compared with myCAFs, csCAFs expressed lower levels of *Tgfb1*, suggesting reduced TGF-β signaling within the tumor microenvironment. This shift was accompanied by decreases in TGF-β response and epithelial-to-mesenchymal transition-associated transcriptional programs within tumor cells, consistent with a less tumor-supportive microenvironment. Collectively, these findings demonstrate that PFA serves not only as a modality for localized tumor destruction but also as a tool to interrogate dynamic tumor-stromal interactions. By enabling controlled, serial perturbation of the pancreatic tumor microenvironment, PFA revealed previously underappreciated plasticity in CAF states and identified stromal remodeling as a potentially important consequence of ablation. These observations establish a framework for using PFA to study and manipulate microenvironmental programs that influence PDAC progression and therapeutic response.

Future studies will be required to evaluate local and distant tumor control and immune responses, as well as rational combinatorial strategies utilizing pulsed field ablation and chemotherapy, immune checkpoint blockade, or other local ablation techniques to leverage these insights into clinically effective interventions for PDAC.

## Acknowledgements

JMB-L received funding from (R01CA277161-01A1), ALPFA, and DOW (HT94252410921). NT and RV report holding stock/equity options in ALPFA, which provided funding for this research NT received funding from R01CA277161-01A1. Research reported in this publication was supported by the National Cancer Institute of the National Institutes of Health under award number P30 CA036727 (UNMC CCSG Cancer Center Grant). The content is solely the responsibility of the authors and does not necessarily represent the official views of the National Institutes of Health.

