## Supplemental Figure 1 for "Pulsed Field Ablation Enables Stromal Manipulation and Tumor Reduction in Pancreatic Ductal Adenocarcinoma"

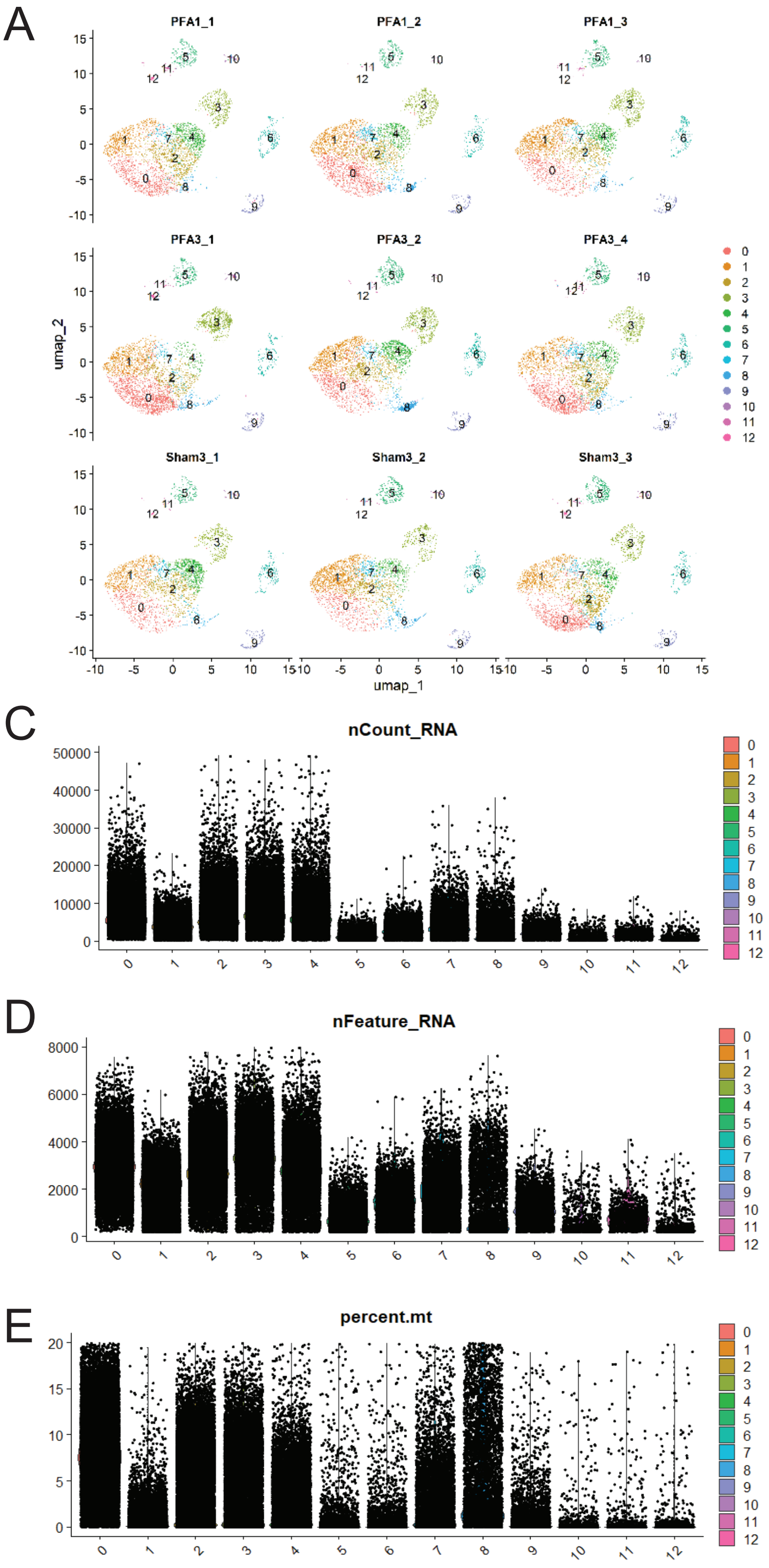


B

| p_val | avg_log2FC | pct.1 | pct.2 | p_val_adj | Cluster | Score | Gene |
|---|---|---|---|---|---|---|---|
| 0 | 5.257322 | 0.668 | 0.16 | 0 | 8 | 2.670719 | Rpl27 |
| 0 | 5.293707 | 0.709 | 0.205 | 0 | 8 | 2.668028 | Rpl34 |
| 0 | 5.128255 | 0.695 | 0.201 | 0 | 8 | 2.533358 | Rps16 |
| 0 | 4.947802 | 0.768 | 0.259 | 0 | 8 | 2.518431 | Rps10 |
| 0 | 5.138739 | 0.667 | 0.177 | 0 | 8 | 2.517982 | Rpl32 |
| 0 | 4.873375 | 0.787 | 0.272 | 0 | 8 | 2.509788 | Tmsb10 |
| 0 | 4.959762 | 0.71 | 0.204 | 0 | 8 | 2.509639 | Rpl14 |
| 0 | 4.620415 | 0.775 | 0.234 | 0 | 8 | 2.499644 | H2az1 |
| 0 | 4.9884 | 0.739 | 0.24 | 0 | 8 | 2.489212 | Rps12 |
| 0 | 5.128653 | 0.697 | 0.212 | 0 | 8 | 2.487397 | Rps15a |
| 0 | 4.894718 | 0.77 | 0.268 | 0 | 8 | 2.457148 | Rps7 |
| 0 | 4.682232 | 0.851 | 0.327 | 0 | 8 | 2.45349 | Rplp1 |
| 0 | 4.824021 | 0.772 | 0.267 | 0 | 8 | 2.436131 | Rpl23a |
| 0 | 4.961667 | 0.679 | 0.192 | 0 | 8 | 2.416332 | Rps11 |
| 0 | 4.646952 | 0.826 | 0.312 | 0 | 8 | 2.388533 | Rps3a1 |
| 0 | 4.914378 | 0.686 | 0.205 | 0 | 8 | 2.363816 | Rps23 |
| 0 | 4.874566 | 0.779 | 0.295 | 0 | 8 | 2.35929 | Rps24 |
| 0 | 5.082068 | 0.622 | 0.163 | 0 | 8 | 2.332669 | Rpl38 |
| 0 | 4.79959 | 0.692 | 0.21 | 0 | 8 | 2.313402 | Rpl15 |
| 0 | 4.76731 | 0.716 | 0.233 | 0 | 8 | 2.302611 | Rpl7 |
| 0 | 4.57674 | 0.819 | 0.319 | 0 | 8 | 2.28837 | Ppia |
| 0 | 5.231603 | 0.572 | 0.135 | 0 | 8 | 2.286211 | Rpl26 |
| 0 | 4.925403 | 0.676 | 0.213 | 0 | 8 | 2.280461 | Rpl37 |
| 0 | 4.707664 | 0.726 | 0.242 | 0 | 8 | 2.278509 | Rps4x |
| 0 | 4.50818 | 0.829 | 0.325 | 0 | 8 | 2.272123 | Rpl13 |
| 0 | 4.516199 | 0.834 | 0.334 | 0 | 8 | 2.2581 | S100a11 |
| 0 | 4.679397 | 0.74 | 0.26 | 0 | 8 | 2.24611 | Rps26 |
| 0 | 4.798428 | 0.69 | 0.222 | 0 | 8 | 2.245664 | Rps28 |
| 0 | 4.888865 | 0.652 | 0.193 | 0 | 8 | 2.243989 | Rps19 |
| 0 | 4.346491 | 0.903 | 0.39 | 0 | 8 | 2.22975 | Rps2 |
| 0 | 4.767175 | 0.657 | 0.191 | 0 | 8 | 2.221504 | Pfn1 |
| 0 | 5.052166 | 0.611 | 0.172 | 0 | 8 | 2.217901 | Rpl35a |
| 0 | 4.581422 | 0.767 | 0.283 | 0 | 8 | 2.217408 | Rpsa |
| 0 | 4.648936 | 0.735 | 0.26 | 0 | 8 | 2.208245 | Rpl41 |
| 0 | 4.526268 | 0.805 | 0.318 | 0 | 8 | 2.204292 | Rpl23 |
| 0 | 4.97753 | 0.61 | 0.169 | 0 | 8 | 2.195091 | Rps5 |
| 0 | 4.545931 | 0.772 | 0.29 | 0 | 8 | 2.191139 | Rpl18a |
| 0 | 4.681332 | 0.731 | 0.263 | 0 | 8 | 2.190863 | Rpl13a |
| 0 | 4.140384 | 0.889 | 0.36 | 0 | 8 | 2.190263 | Actb |
| 0 | 4.765265 | 0.681 | 0.222 | 0 | 8 | 2.187257 | Rpl21 |
| 0 | 4.567077 | 0.73 | 0.257 | 0 | 8 | 2.160228 | Rplp0 |
| 0 | 4.58782 | 0.721 | 0.251 | 0 | 8 | 2.156275 | Rpl19 |
| 0 | 4.761853 | 0.649 | 0.199 | 0 | 8 | 2.142834 | Rps3 |
| 0 | 4.578303 | 0.696 | 0.229 | 0 | 8 | 2.138067 | Rpl8 |
| 0 | 4.9036 | 0.612 | 0.176 | 0 | 8 | 2.137969 | Rpl28 |
| 0 | 4.508123 | 0.781 | 0.307 | 0 | 8 | 2.13685 | Fau |
| 0 | 4.675751 | 0.692 | 0.237 | 0 | 8 | 2.127467 | Rpl37a |
| 0 | 4.449461 | 0.699 | 0.223 | 0 | 8 | 2.117943 | Gapdh |
| 0 | 4.402791 | 0.829 | 0.348 | 0 | 8 | 2.117742 | Rps8 |

**Supplemental Figure 1. Cluster 8 displays low-quality characteristics. A.** Distribution of clusters between samples. **B**. Top differentially expressed genes in cluster 8 is dominated by ribosomal genes. **C**. RNA count across clusters. **D**. RNA features across samples. **E**. Percent of reads that are mitochondrial genes across clusters.
