## Supplemental Figure 2 for "Pulsed Field Ablation Enables Stromal Manipulation and Tumor Reduction in Pancreatic Ductal Adenocarcinoma"

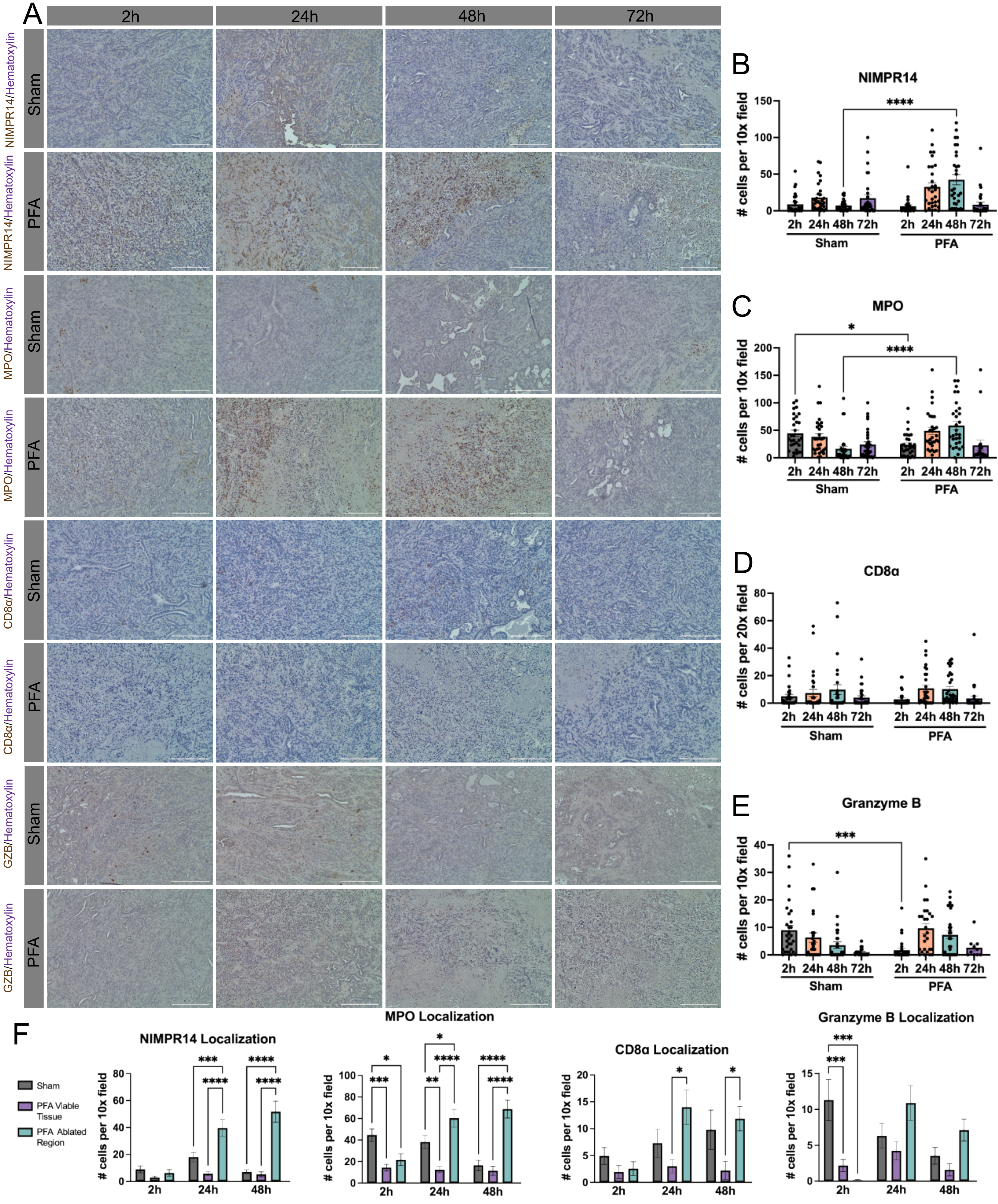


**Supplemental Figure 2. PFA treatment induces early neutrophil infiltration. A.** Representative IHC images for NIMPR14, MPO, CD8α, and Granzyme B at each time point. **B.** NIMPR14 is significantly increased in PFA 48 hours post ablation. **C.** MPO is significantly increased in Sham 2 hours post ablation and significantly increased in PFA 48 hours post ablation. **D.** There are no significant differences in CD8α quantification. **E.** Granzyme B is significantly higher in Sham at 2 hours post ablation. **F.** NIMPR14, MPO, and CD8α show significant localization to the ablated region within PFA-treated tumors. Data presented as mean ± SEM. *p<0.05. **p<0.01, ***p<0.001, ****p<0.0001.
