## Supplemental Figure 3 for "Pulsed Field Ablation Enables Stromal Manipulation and Tumor Reduction in Pancreatic Ductal Adenocarcinoma"

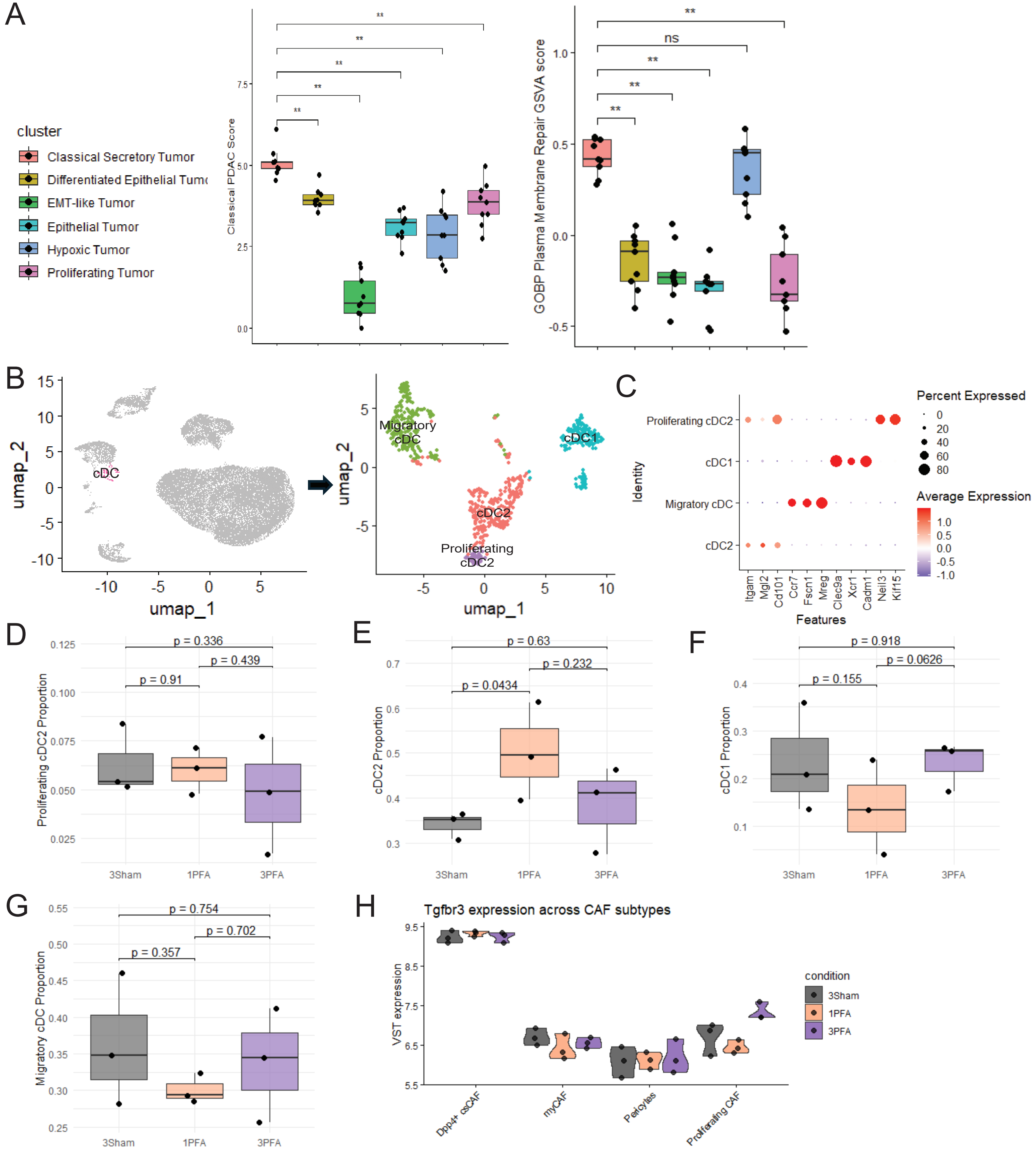


**Supplemental Figure 3. Analysis of tumor, immune, and CAF clusters. A.** Pathways that show enrichment in the classical secretory tumor cluster. **B.** cDC sub-clustering reveals four distinct subtypes. **C.** Top differentially expressed markers for each cDC subtype. **D.** Proportion of cDC1 is consistent across groups. **E.** Proportion of cDC2 is elevated in 1 PFA when compared to 3 Sham. **F.** Proportion of migratory cDC is consistent across groups. **G.** Proportion of proliferating cDC2 is consistent across groups. **H.** Variance-stabilized transformation Tgfbr3 expression across CAF subtypes.
