## Supplemental Figure 4 for "Pulsed Field Ablation Enables Stromal Manipulation and Tumor Reduction in Pancreatic Ductal Adenocarcinoma"

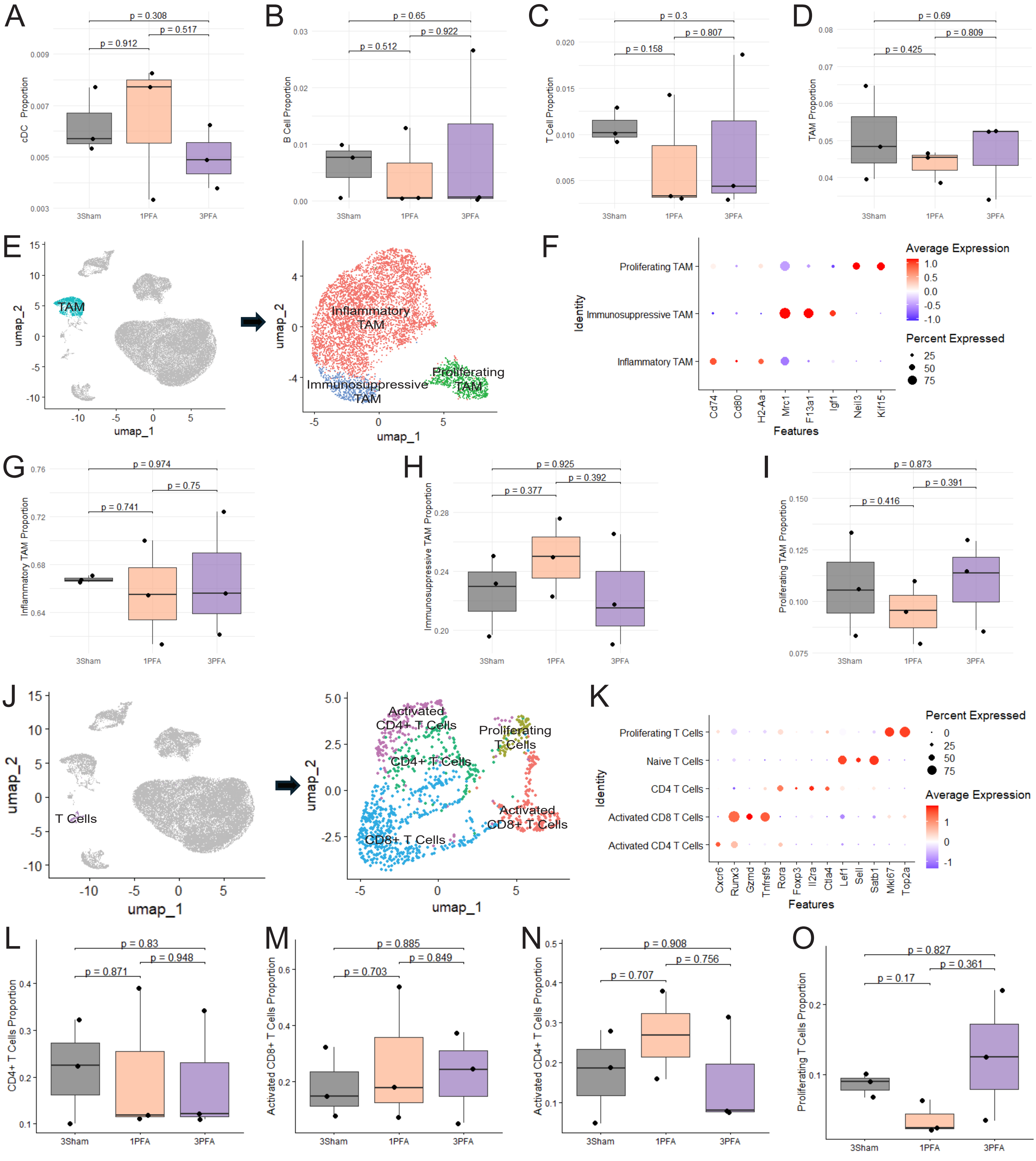


**Supplemental Figure 4. PFA does not induce changes in the phenotype of either macrophages or T cells. A.** Proportion of classic dendritic cells is consistent across groups. **B.** Proportion of B cells is consistent across groups. **C**. Proportion of T cells is consistent across groups. **D.** Proportion of tumor associated macrophages (TAM) is consistent across groups. **E.** TAM sub-clustering reveals three distinct subtypes. **F.** Top differentially expressed markers for each TAM subtype. **G.** Proportion of inflammatory TAMs is consistent across groups. **H.** Proportion of immunosuppressive TAMs is consistent across groups. **I.** Proportion of proliferating TAMs is consistent across groups. **J.** T cell sub-clustering reveals three distinct subtypes. **K.** Top differentially expressed markers for each T cell subtype. **L.** Proportion of activated CD4+ T cells is consistent across groups. **M.** Proportion of activated CD8+ T cells is consistent across groups. **N.** Proportion CD4+ T cells is consistent across groups. **O.** Proportion of proliferating T cells is consistent across groups.
