## Supplemental Figure 5 for "Pulsed Field Ablation Enables Stromal Manipulation and Tumor Reduction in Pancreatic Ductal Adenocarcinoma"

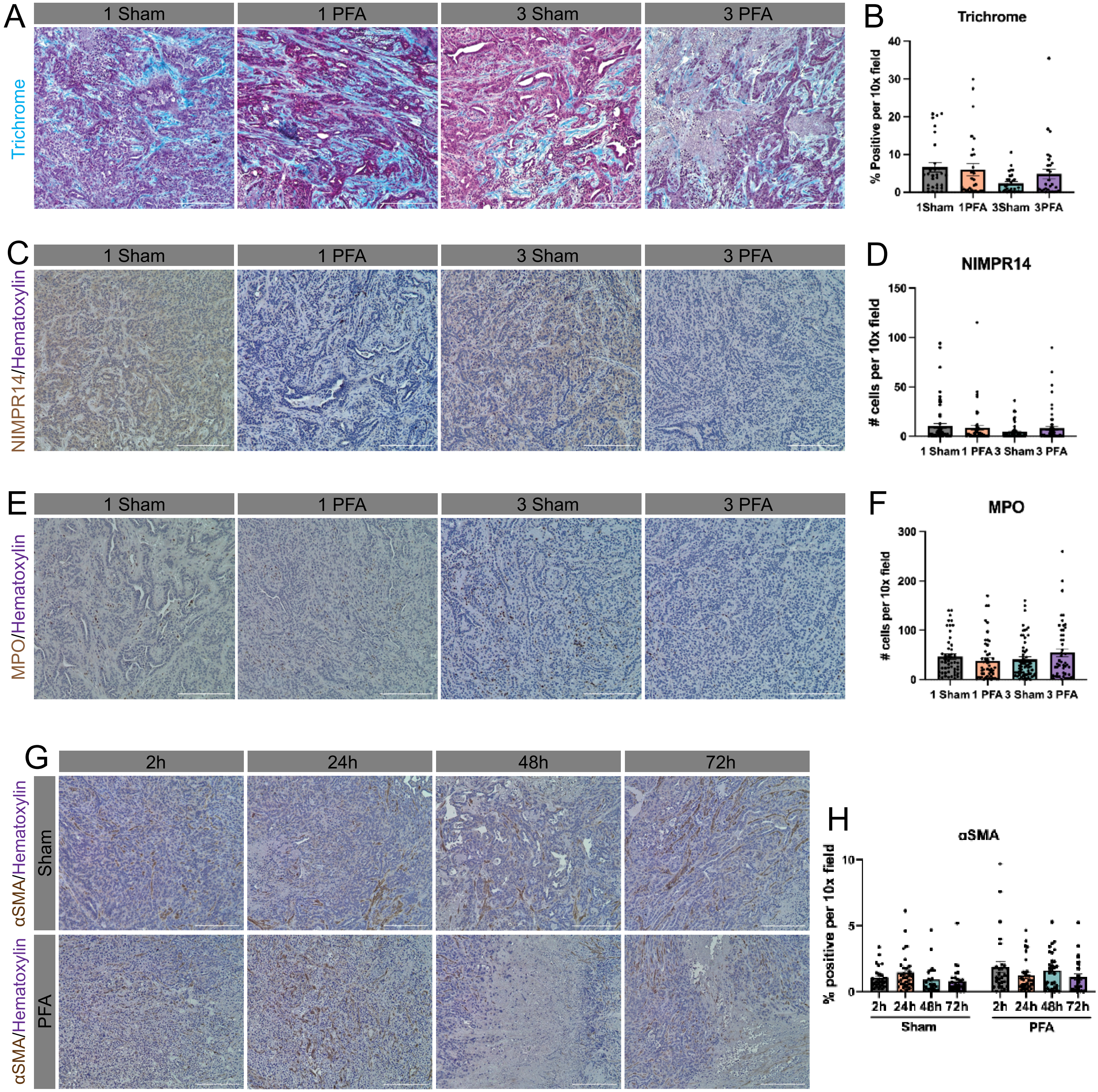


**Supplemental Figure 5. Tumor characterization across experiments. A.** Representative trichrome images at each time point. **B**. There is no significant difference in trichrome staining between groups. **C**. Representative IHC images of NIMPR14 at each time point. **D**. There are no significant differences in NIMPR14 staining between groups. **E**. Representative IHC images of MPO after multiple PFA sessions. **F**. There are no significant differences in MPO staining between groups. **G**. Representative IHC images of αSMA after multiple PFA sessions. **H**. There are no significant differences in αSMA staining between groups. Scale bars are 25 μm. Data presented as mean ± SEM.
