## Supplemental Table 1 for "Pulsed Field Ablation Enables Stromal Manipulation and Tumor Reduction in Pancreatic Ductal Adenocarcinoma"

**Immunohistochemistry antibody dilution**

| Antibody | Catalogue # | Dilution | Antigen Retrivel pH | Secondary Antibody Host | Secondary Dilution |
|---|---|---|---|---|---|
| NIMPR14 | ab2557 | 1:100 | pH 9 | Rat | 1:500 |
| MPO | ab208670 | 1:300 | pH 9 | Rabbit | 1:500 |
| CD8α | ab217344 | 1:250 | pH 9 | Rabbit | 1:500 |
| GZMB | ab255598 | 1:200 | pH 9 | Rabbit | 1:500 |
| αSMA | ab5694 | 1:500 | pH 6 | Rabbit | 1:500 |
