## Supplemental Table 2 for "Pulsed Field Ablation Enables Stromal Manipulation and Tumor Reduction in Pancreatic Ductal Adenocarcinoma"

| Sample | Raw Cells | Post-QC | QC Remov | % QC Rem | Dbls Remo | % Dbls Rer | Post Doubl | Post Low-Quality Clust | |
|---|---|---|---|---|---|---|---|---|---|
| PFA1_1 | 19267 | 18904 | 363 | 1.88405 | 3242 | 17.14981 | 15662 | 15258 | |
| PFA1_2 | 17981 | 17625 | 356 | 1.979868 | 2193 | 12.44255 | 15432 | 14658 | |
| PFA1_3 | 16251 | 16120 | 131 | 0.806104 | 1969 | 12.21464 | 14151 | 13596 | |
| PFA3_1 | 21504 | 20769 | 735 | 3.417969 | 3633 | 17.49242 | 17136 | 16335 | |
| PFA3_2 | 19870 | 19576 | 294 | 1.479618 | 2332 | 11.91255 | 17244 | 15319 | |
| PFA3_4 | 20483 | 19744 | 739 | 3.60787 | 3157 | 15.98967 | 16587 | 15974 | |
| Sham3_1 | 15466 | 15287 | 179 | 1.157377 | 1790 | 11.7093 | 13497 | 12996 | |
| Sham3_2 | 16725 | 16388 | 337 | 2.014948 | 2305 | 14.06517 | 14083 | 13472 | |
| Sham3_3 | 19848 | 18748 | 1100 | 5.54212 | 2931 | 15.63367 | 15817 | 15183 | |
| Total: | 167395 | | | | | | Total post ( | 132791 | 79.32794 |

:er Removal (Final Cells)

% retained
