## Supplemental Table 3 for "Pulsed Field Ablation Enables Stromal Manipulation and Tumor Reduction in Pancreatic Ductal Adenocarcinoma"

| Package | Version | Function | Citation |
| --- | --- | --- | --- |
| ComplexHeatmap | v2.26.1 | Heatmap generation | (6) |
| ggplot2 | v4.0.3 | Generate box plots and violin plots | (7) |
| speckle | v1.10.0 | Analyze difference in cell proportion | (8) |
| DESeq2 | v1.50.2 | Estimates fold change and dispersion of expression | (9) |
| msigdbr | v26.1.0 | Used to access gsea libraries | (10) |
| edgeR | v4.8.2 | Generates differentially expressed genes for gsea | (11) |
| fgsea | v1.36.2 | Performs fast gsea analysis | (12) |
| enrichplot | v1.30.5 | Generates running enrichment plots | (13) |
| GSVA | v2.4.9 | Generates gene set variation analysis | (14) |
